# CB2 distinguishes cells from background barcodes in 10x Genomics data

**DOI:** 10.1101/832535

**Authors:** Zijian Ni, Shuyang Chen, Jared Brown, Christina Kendziorski

**Affiliations:** Department of Statistics, University of Wisconsin-Madison, Madison, WI, USA; Department of Biostatistics and Medical Informatics, University of Wisconsin-Madison, Madison, WI, USA

## Abstract

An important challenge in pre-processing data from the 10x Genomics Chromium platform is distinguishing barcodes associated with real cells from those binding background reads. Existing methods test barcodes individually, and consequently do not leverage the strong cell-to-cell correlation present in most datasets. To improve the power to identify real cells and rare subpopulations, we introduce CB2, a cluster-based approach for distinguishing real cells from background barcodes.

---

The 10x Genomics Chromium (10x)^1^ platform is a powerful and widely-used approach for profiling genome-wide gene expression in individual cells. The technology utilizes gel beads, each containing oligonucleotide indexes made up of bead-specific barcodes combined with unique molecular identifiers (UMIs)^2^ and oligo-dT tags to prime polyadenylated RNA. Single cells of interest are combined with reagents in one channel of a microfluidic chip, and gel beads in another, to form gel-beads in emulsion, or GEMs. Oligonucleotide indexes bind polyadenylated RNA within each GEM reaction vesicle before gel beads are dissolved releasing the bound oligos into solution for reverse transcription. By design, each resulting cDNA molecule contains a UMI and a GEM-specific barcode. Indexed cDNA is pooled for PCR amplification and sequencing resulting in a data matrix of UMI counts for each barcode (**Supplementary Figure 1**).

Ideally, each barcode will tag mRNA from an individual cell, but this is often not the case in practice. In most datasets, approximately 90% of GEMs do not contain viable cells, but rather contain ambient RNA excreted by cells in solution or as a product of cell lysis^1^. As a result, an important challenge in pre-processing 10x data is distinguishing those barcodes corresponding to real cells from those binding ambient, or background, RNA.

Early methods to address this challenge defined real cells as those barcodes with total read counts exceeding some threshold^1,3^. Such methods are suboptimal as they discard small cells as well as those expressing relatively few genes. To address this, Lun *et al*., 2019^4^ developed EmptyDrops (ED), an approach to identify individual barcodes with distributions varying from a background distribution. Similar to previous approaches, ED identifies an upper threshold and defines real cells as those barcodes with counts above the threshold. As a second step, ED uses all barcodes with counts below a lower threshold to estimate a background distribution of ambient RNA against which remaining barcodes are tested. Those having expression profiles significantly different from the background distribution are deemed real cells. The ED approach is current state-of-the-art in the field. However, given that ED performs tests for each barcode individually, it does not leverage the strong correlation observed between cells and, consequently, compromises power in many datasets.

To increase the power for identifying real cells, we propose CB2, a <u>c</u>luster-<u>b</u>ased approach for distinguishing real <u>c</u>ells from <u>b</u>ackground barcodes in 10x experiments. CB2 extends the ED framework by introducing a clustering step that groups similar barcodes, then conducts a statistical test to identify groups with expression distributions that vary from the background (**Figure 1, Supplementary Figure 2**). CB2 is implemented in the R package *scCB2*.

**Figure 1:**
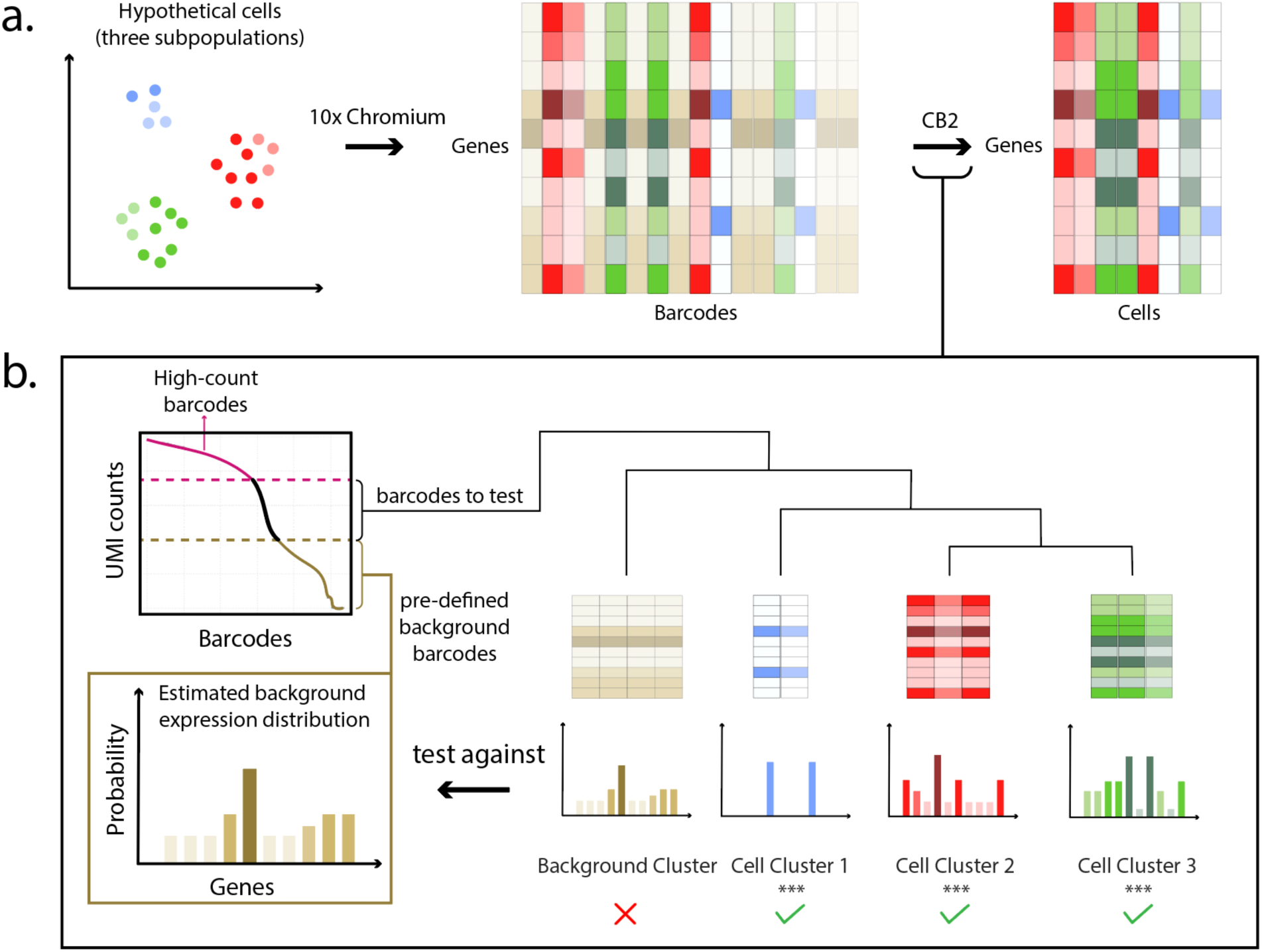
Overview of CB2. (a) Projection of a hypothetical cell population containing three sub-populations (red, green, and blue where intensity corresponds to read depth). CB2 takes as input a gene by barcode matrix of UMI counts and returns a gene by cell matrix. (b) High-count barcodes with counts above a pre-specified upper threshold are considered real cells; barcodes with counts below a lower threshold are used to estimate a background distribution (**Supplementary Figure 2**). The remaining barcodes are clustered and tight clusters are tested as a group against the estimated background distribution; barcodes not in tight clusters are tested individually (not shown). High-count barcodes and those identified by CB2 are retained for downstream analysis.

CB2 was evaluated and compared with ED on simulated and case study data. In SIM IA, counts are generated as in Lun *et al*., 2019^4^. Briefly, given an input dataset, an inflection point dividing low from high count barcodes is determined. Low count barcodes are pooled to estimate the background distribution. Two thousand barcodes are then randomly sampled from the high count barcodes to give real cells (referred to as *G*_1_ cells^4^); a second set of 2000 high count barcodes is sampled and then downsampled by 90% to give *G*_2_ cells. We added a third set (*G*_1.5_) by sampling 2000 barcodes from the high count range and downsampling by 50%. **Supplementary Figure 3** shows increased power of CB2 with well controlled false discovery rate (FDR) for the 6 datasets considered in Lun *et al*., 2019^4^ as well as 4 additional datasets. SIM IB, also considered by Lun *et al*., 2019^4^, is similar to SIM IA, but in SIM IB 10% of the genes in the real cells are shuffled making the real cells more different from the background and therefore easier to identify (**Supplementary Figure 4**). **Supplementary Figure 5** shows the increased power of CB2 is maintained.

To further evaluate CB2, we applied CB2 and ED to the ten datasets evaluated in the simulation study as well as one additional dataset considered in the ED case study and compared the number of cells identified in common as well as those uniquely identified by each approach.

**Supplementary Table 1** shows that CB2 finds 24% more cells on average (range 4%-81%). Of the extra cells identified, 88% on average (range 44%-100%) add to existing subpopulations. The remaining 12% (range 0%-56%) make up novel subpopulations. As an example, **Figure 2** and **Supplementary Figure 6** show results from the Alzheimer data^5^ where CB2 identifies 18% more cells within excitatory neuron, inhibitory neuron, and oligodendrocyte sub-populations, and also reveals a novel subpopulation consisting of 209 cells. The novel subpopulation uniquely shows high expression of both oligodendrocyte and astrocyte marker genes, suggesting that this group may be mixed phenotype glial cells^6^ (**Supplementary Figure 6**). By increasing the number of true cells identified, CB2 also improves the power to differentiate Alzheimer’s patients from controls (**Supplementary Figure 7**).

**Figure 2:**
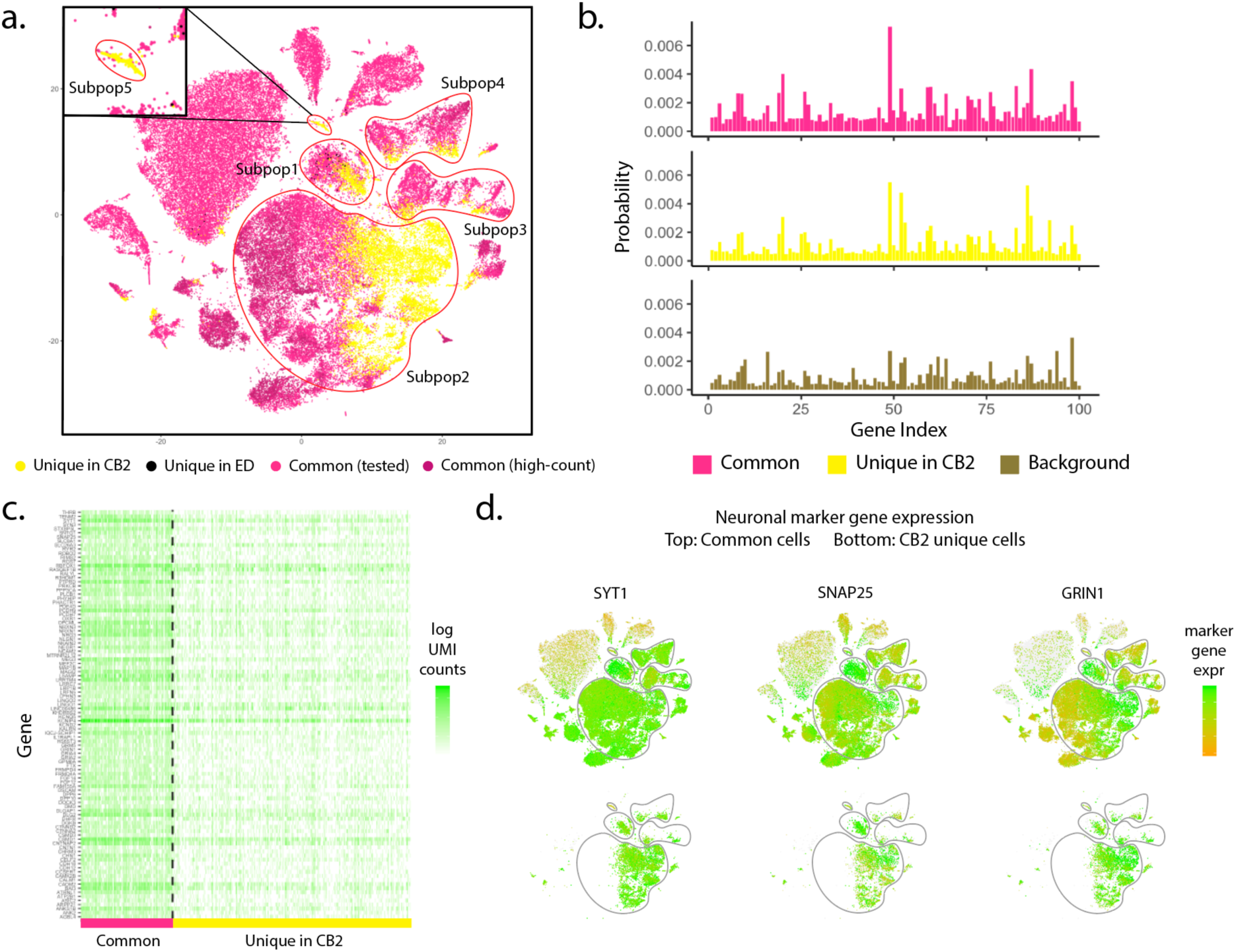
Results from the Alzheimer dataset. (a) t-SNE plot of cells identified by CB2 and ED. High-count barcodes exceeding an upper threshold are identified as real cells by both methods without a statistical test (dark pink); barcodes identified as cells by both methods following statistical test are shown in pink. Cells identified uniquely by CB2 (yellow) and ED (black) are also shown. CB2 identifies an increased number of cells in existing sub-populations (Subpop1 – Subpop4) and also identifies a novel subpopulation (Subpop5). (b) Distribution plots of the 100 genes having highest average expression in Subpop2 are shown for cells identified by both CB2 and ED (upper) and identified uniquely by CB2 (middle). The estimated background distribution is also shown (lower). Cells uniquely identified by CB2 in Subpop2 have a distribution similar to other Subpop2 cells and differ from the background. (c) Heatmap of log transformed raw UMI counts for the same 100 genes for barcodes identified by CB2 and ED (left) and barcodes uniquely identified by CB2 (right). (d) t-SNE plots of cells colored by neuron marker genes SYT1, SNAP25, and GRIN1 in all cells (upper) and those identified uniquely by CB2 (lower).

Results from three additional datasets are shown in **Supplementary Figures 8-10. Figure 2** and **Supplementary Figures 6-10** demonstrate that CB2 provides a powerful approach for distinguishing real cells from background barcodes which will increase the number of cells identified in existing cell subpopulations in most datasets and may facilitate the identification of novel subpopulations.

The 10x Genomics Chromium platform provides unprecedented opportunity to address biological questions, but efficient pre-processing is required to maximize the information obtained in an experiment. Our approach allows investigators to maximize the number of cells retained, and consequently to increase the power and precision of downstream analysis.

## ACCESSION CODES

The Alzheimer case study dataset was downloaded from https://www.synapse.org/#!Synapse:syn16780177^5^. The placenta dataset^7^ is available at https://jmlab-gitlab.cruk.cam.ac.uk/publications/EmptyDrops2017-DataFiles. All other datasets in this study are available at the 10x Genomics website (https://support.10xgenomics.com/single-cell-gene-expression/datasets) (**Supplementary Table 2**). The R package R/*scCB2* is available at https://github.com/zijianni/scCB2, and will be submitted to Bioconductor^8^. All simulation codes and a case study data analysis script are available at https://github.com/zijianni/codes-for-CB2-paper.

## ACKNOWLEDGMENTS

This work was supported by NIH GM102756. The authors thank Matt Bernstein, Chitrasen Mohanty, and Ziyue Wang for comments that improved the manuscript.

## AUTHOR CONTRIBUTIONS

Z.N. and C.K. designed the research, developed the method, and wrote the first version of the manuscript. Z.N. and S.C. conducted simulations and quality control evaluations. Z.N., S.C., and C.K. built and tested the R package. Z.N., J.B., and C.K. analyzed results from early versions of the method which helped during method refinement. All authors contributed to writing the manuscript.

## ONLINE METHODS

### Versions

For cell identification with R/*scCB2* 0.99.12 and R/*DropletUtils* 1.5.4^4,9^, the latest version of R^10^ was used: 3.7-devel (2019-07-17 r76847). Other packages are not yet compatible or not stable with the R developers version and so for *scran* 1.12.1^11^, *Seurat* 3.1.0^12,13^, and *ggplot2* 3.2.1^14^, R 3.6.0 (2019-04-24 r76423) was used.

### CB2

As CB2 relies on ED, we briefly review the ED approach before detailing the clustering test introduced in CB2. ED expects as input a *G* × *B* feature-by-barcode matrix with *G* features (for simplicity, we refer to features as genes) and *B* barcodes. Barcodes having zero counts for all genes are filtered out and the remaining barcodes are divided into three groups based on the sum of gene expression (UMI) counts within a barcode. The background group, *B*_0_, contains all barcodes with counts less than or equal to a pre-defined lower threshold (defaults to 100); the high-count barcodes, *B*_2_, contain barcodes with counts exceeding an upper threshold (defaults to knee point); the remaining barcodes (*B*_1_) are tested (**Supplementary Figure 2**).

ED assumes that counts from a background barcode are distributed as Dirichlet-Multinomial with probability vector 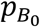 estimated by averaging the counts in *B*_0_ and applying the Good-Turing algorithm^15^ to ensure that all probabilities are non-zero, denoted as 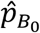. For a barcode *b* ∈ *B*_1_, ED tests 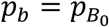 against the alternative 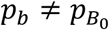 using the log-likelihood under 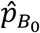 as the test statistic. A Monte-Carlo p-value is calculated via simulating Dirichlet-Multinomial barcodes of size |*b*| under 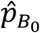 and calculating the proportion of simulated barcodes having a test statistic more extreme than (or equal to) *b*’s. The false discovery rate is controlled using the Benjamini-Hochberg procedure^16^.

CB2 follows ED by filtering out genes with zero counts and dividing the remaining barcodes into three groups. However, instead of testing all barcodes from *B*_1_ individually, CB2 first clusters barcodes and then tests tight clusters to identify those that differ from the background. CB2 proceeds as follows:

1. **Barcodes grouped by size.** CB2 orders barcodes in *B*_1_ by total counts

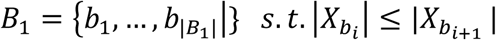

where *X*_*b*_ denotes the count vector of barcode *b*, |*X*_*b*_| denotes the total UMI count of barcode *b*, and |*B*_1_| denotes the number of barcodes in *B*_1_. Groups of size *S* (defaults to 1000 in R*/scCB2*) are constructed consisting of barcodes ranging in size from smallest to largest:

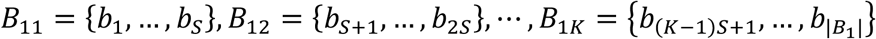

where 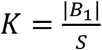 is rounded up if not an integer. If 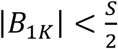, barcodes in *B*_1*K*_ are merged with those in *B*_1(*K*−1)_. Sorting barcodes by size reduces bias in the clustering and testing steps that follow.
2. **Barcodes clustered within group:** Barcodes within each group *B*_1*j*_ are clustered using hierarchical clustering with pairwise Pearson correlation as the similarity metric. A cluster is considered tight if the average within-cluster pairwise Pearson correlation exceeds a data-driven threshold. Tight clusters are retained for further analysis as described in step 3, below. To determine thresholds, ten tight clusters of varying size are simulated by generating 100 samples from a Multinomial distribution with parameters (*N, p*) where *N* ranges from 100 to 1000 in increments of size 100. This range is chosen as we found little variation in thresholds for barcode sizes exceeding 1000; *p* is set to either 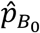 or 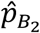, whichever has larger Shannon entropy^17^ as the distribution with larger entropy is less affected by outliers. For each simulated cluster *C*, the threshold *κ*_*c*_ is defined by its average pairwise Pearson correlation. A cluster is considered tight if the average within-cluster pairwise Pearson correlation exceeds *κ*_*c*_ for the simulated cluster of closest size.
3. **Tight clusters tested:** For each tight cluster *C*, we conduct a Monte-Carlo test to assess dissimilarity from the background. Pairwise Pearson correlations are calculated between every barcode in *C* and 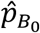; the test statistic for cluster *C, T*_*c*_, is defined to be the median of these correlations. Similar to ED, to simulate background barcodes, we sample barcodes 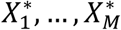 from a Multinomial 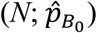 where *N* is the size of the barcode giving *T*_*c*_. The Monte-Carlo p-value is:

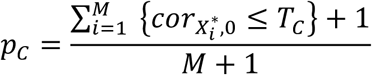

where 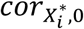 is the Pearson correlation between 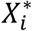 and 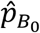 (*M* defaults to 1000 in R/*scCB2*). Monte-Carlo p-values are calculated for each cluster followed by Benjamini-Hochberg^16^ to control the FDR. All barcodes within a significant cluster are identified as real cells.
4. **Individual barcodes tested:** Barcodes that were not included in a tight cluster in Step 2 as well as those in a tight cluster that were not found to be significant in Step 3 are tested individually using ED. It is important to note that some of the barcodes identified in this step do not overlap with identifications made when ED is applied to the full set of barcodes given differences in the rates of true cells to background barcodes and differences in error rate control.

### Simulations

Counts are generated as in Lun *et al.* 2019^4^. As detailed there, each simulation requires an input dataset. We constructed simulations from 10 datasets: Alzheimer^5^, PBMC8K, PBMC33K, mbrain1K, mbrain9K, PanT4K, MALT, PBMC4K, jurkat, and T293 (**Supplementary Table 2**). For each input dataset, the inflection point of the UMI count by sorted barcode plot is used to divide lower count from higher count barcodes. The barcodes in the lower count range are considered background. In SIM IA, two sets of 2000 barcodes randomly sampled from the higher count range are considered real cells. The first set of 2000 is referred to as large (*G*_1_) cells; the second set is downsampled by 90% to give small (*G*_2_) cells. We added a third set of medium (*G*_1.5_) cells by sampling 2000 cells from the higher count range and downsampling by 50%. The process for simulating data in SIM IB is identical to SIM IA except that in SIM IB, 10% of the genes in each simulated real cell are shuffled making the real cells more different from the background barcodes and, consequently, making real cells easier to identify. SIM IA is a more realistic simulation (**Supplementary Figure 4**).

### Case studies

We evaluated the 10 datasets used in the simulation and also the placenta data evaluated in Lun *et al.* 2019^4^. These datasets vary in sequencing depth as well as in the extent of differences between the real cell and background distributions (**Supplementary Figure 4**). CB2 and ED were applied to each dataset using default settings. For plots that compare identifications between CB2 and ED, cells identified by either approach (or both) were combined and UMI counts were normalized via *scran*. The *Seurat* pipeline was used to generate t-SNE plots from the top 4000 most highly variable genes and top 50 principal components. Expression heatmaps show log transformed raw UMI counts. For heatmaps and distribution plots, mitochondrial and ribosomal genes were removed.

### Differential expression analysis in Alzheimer data

Cells identified by CB2, ED, or both were combined into a single matrix and filtered similar to Mathys *et al*.^5^. Specifically, cells with mitochondrial gene expression making up 40% or more of the total UMI counts were removed; genes detected in fewer than two cells were also excluded giving a matrix of 28208 genes and 74579 barcodes. Normalization was performed using *scran*. Cell types were annotated using marker genes as in Mathys *et al*.^5^ Differential expression (DE) tests between cells from Alzheimer’s cases and controls were conducted using Wilcoxon rank-sum tests as in Mathys *et al*.^5^. Results were compared for known DE genes extracted from Mathys *et al*.^5^.

### Implementation of CB2 and ED

For all simulation and case study analyses, CB2 and ED were implemented using default parameters. A target FDR was set at 1%.

### Existing subpopulations vs. novel subpopulations

The *FindNeighbors* and *FindClusters* functions in *Seurat* were used with default settings to assign each cell to a cluster, referred to here as a subpopulation. For each subpopulation, we calculated the percentage of cells identified by both CB2 and ED as well as those identified uniquely by CB2. Subpopulations for which over 80% of the cells are uniquely identified by CB2 are referred to as novel subpopulations (**Supplementary Table 3** shows the number of novel subpopulations identified using 70%, 80%, or 90% as thresholds).

