## Supplementary Materials for "CB2 distinguishes cells from background barcodes in 10x Genomics data"

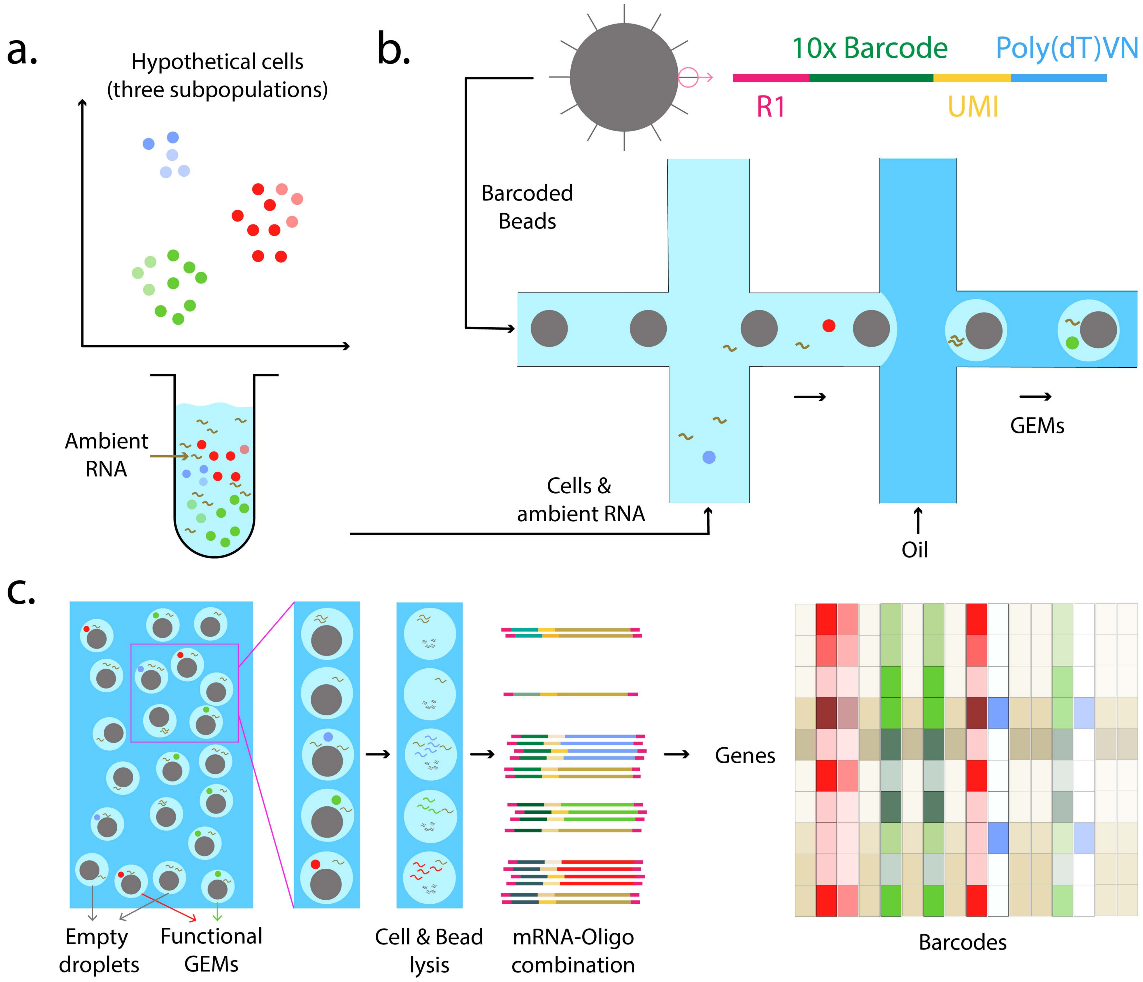


**Supplementary Figure 1:** (a) Projection of a hypothetical cell population containing three sub-populations (red, green and blue where intensity corresponds to read depth). (b) In the 10x RNA-sequencing protocol[1], gel beads containing oligonucleotide indexes made up of bead-specific barcodes combined with UMIs and oligo-dT tags to prime polyadenylated RNA are combined with single cells in one channel of a microfluidic chip and oil in another to form gel-beads in emulsion, or GEMs. (c) GEMs that capture individual cells are referred to as functional GEMs; those that fail to capture cells are empty droplets. Gel beads dissolve and release their oligos for reverse transcription of polyadenylated RNAs. (d) Indexed cDNA is pooled for PCR amplification and sequencing to give a data matrix of UMI counts for each barcode, which is taken as input to CB2 (**Figure 1**). By leveraging correlation among cell sub-populations, CB2 shows improved sensitivity and specificity for identifying cells in common as well as rare subpopulations.

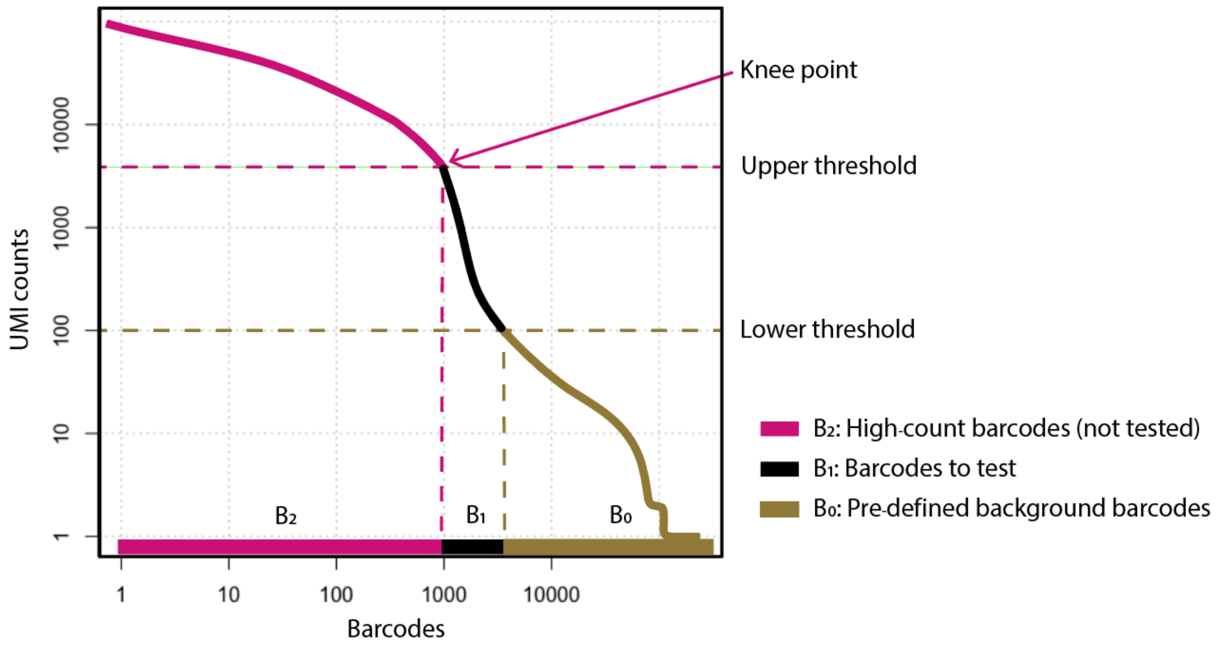


**Supplementary Figure 2**: UMI counts are plotted against barcodes that have been ordered by total count. Both CB2 and ED automatically call high count barcodes real cells (they are not tested); low count barcodes are considered background. The remaining barcodes are tested. By default, the high count (upper) threshold is defined by the knee point, where the counts vs. rank curve begins to drop rapidly; the low count threshold is set to 100. Either, or both, may be changed by a user.

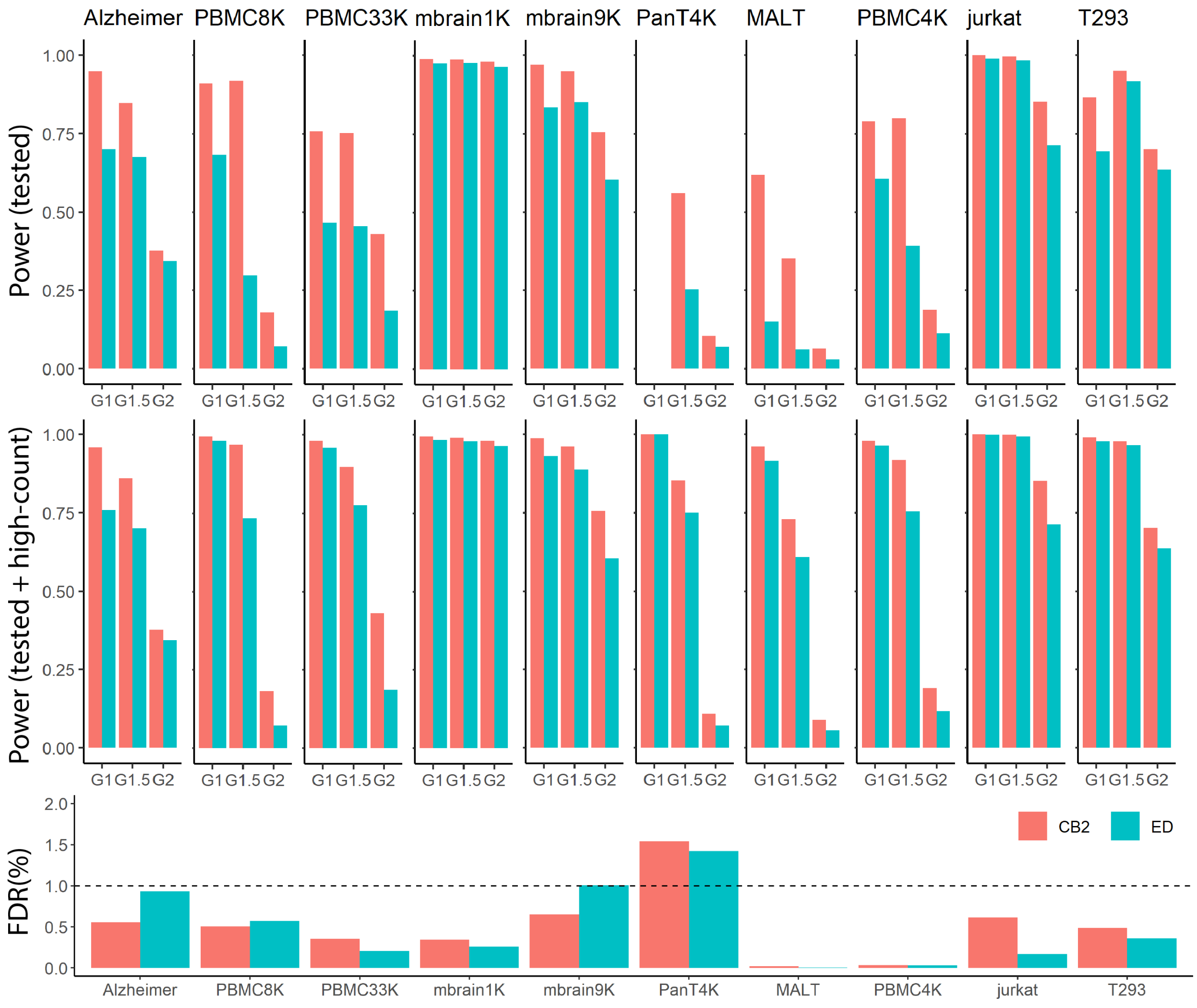


**Supplementary Figure 3:** The average power and average false discovery rate (FDR) of CB2 and ED for SIM IA data (average taken over 5 simulated datasets). Since both CB2 and ED automatically identify high count barcodes as real cells (they are not subject to statistical test; **Supplementary Figure 2**), we report results for all barcodes as well as those tested by CB2 and ED. The top panel shows the average power for tested barcodes; the middle panel for tested as well as high count barcodes. The bottom panel shows the average FDR. For the PanT4K dataset, all $G_1$ cells are above the upper threshold and so no barcodes were tested (as a result, power for tested barcodes is not defined).

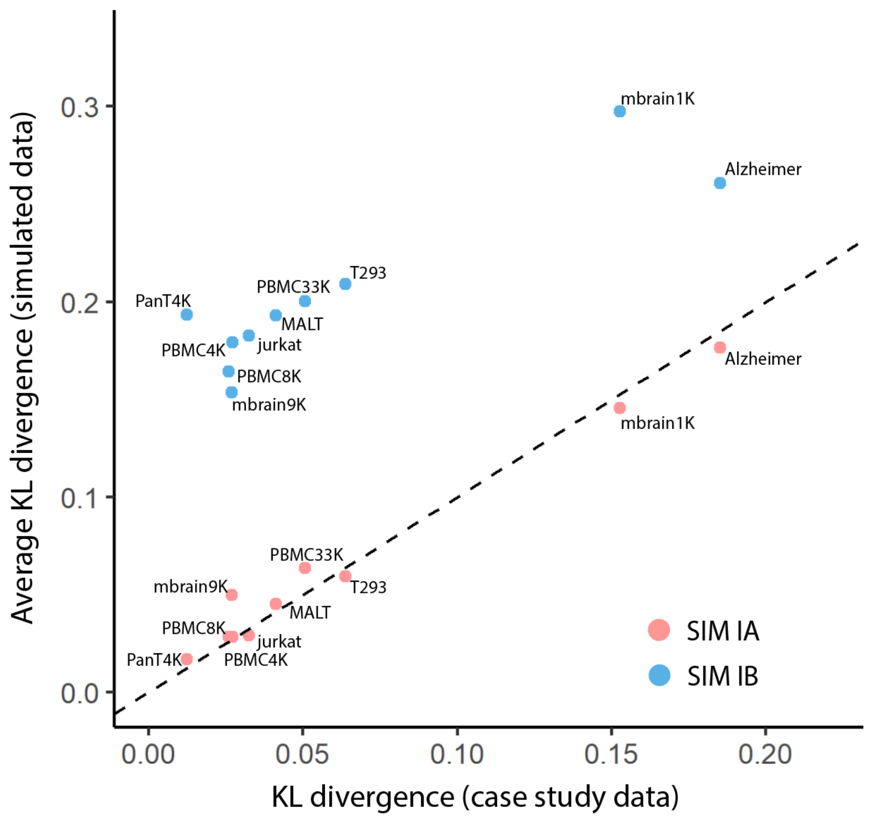


**Supplementary Figure 4:** To measure the similarity between each simulated dataset and the case study data from which it is derived, for each dataset we evaluate the extent to which the distribution of real cells differs from the background distribution. Specifically, for each case study dataset, we calculate the Kullback-Leibler (KL) divergence to measure the difference between the expression distribution of the high-count barcodes and the background distribution. For the simulated data, we calculate the KL divergence between the $G_1$ simulated cells and the background distribution. This is repeated to get a KL divergence for $G_{1.5}$ simulated cells vs. background and $G_2$ simulated cells vs. background. The average KL divergence (averaged over the three groups) for each simulation (SIM IA, SIM IB) is plotted against the KL divergence for each case study. The KL divergence in the SIMIA data is more similar to the case study data for each dataset considered. The increased KL divergence observed in the SIMIB data indicates that the SIMIB simulated barcodes differ from the background more than observed in the case study or SIMIA data, which makes differences easier to identify when applying CB2 or ED.

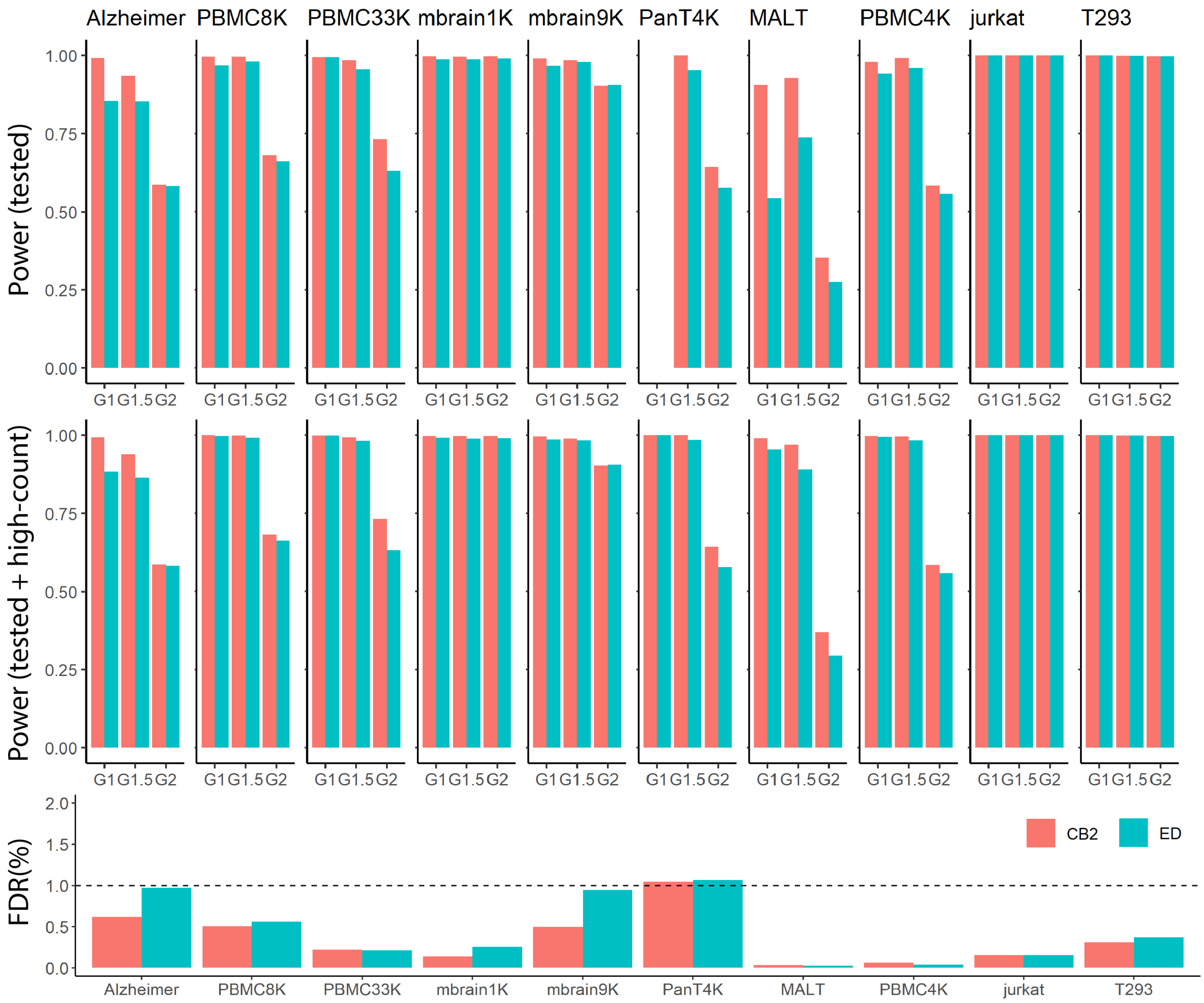


**Supplementary Figure 5:** The average power and average FDR of CB2 and ED for SIM IB data (average taken over 5 simulated datasets). SIM IB, also considered by Lun *et al.*, 2019[2], is similar to SIM IA, but in SIM IB 10% of the genes in the real cells are shuffled making the real cells more different from the background and therefore easier to identify (**Supplementary Figure 4**). Since both CB2 and ED automatically identify high count barcodes as real cells (they are not subject to statistical test; **Supplementary Figure 2**), we report results for all barcodes as well as those tested by CB2 and ED. The top panel shows the average power for tested barcodes; the middle panel for tested as well as high count barcodes. The bottom panel shows the average FDR. For the PanT4K dataset, all $G_1$ cells are above the upper threshold and so no barcodes were tested (as a result, power for tested barcodes is not defined).

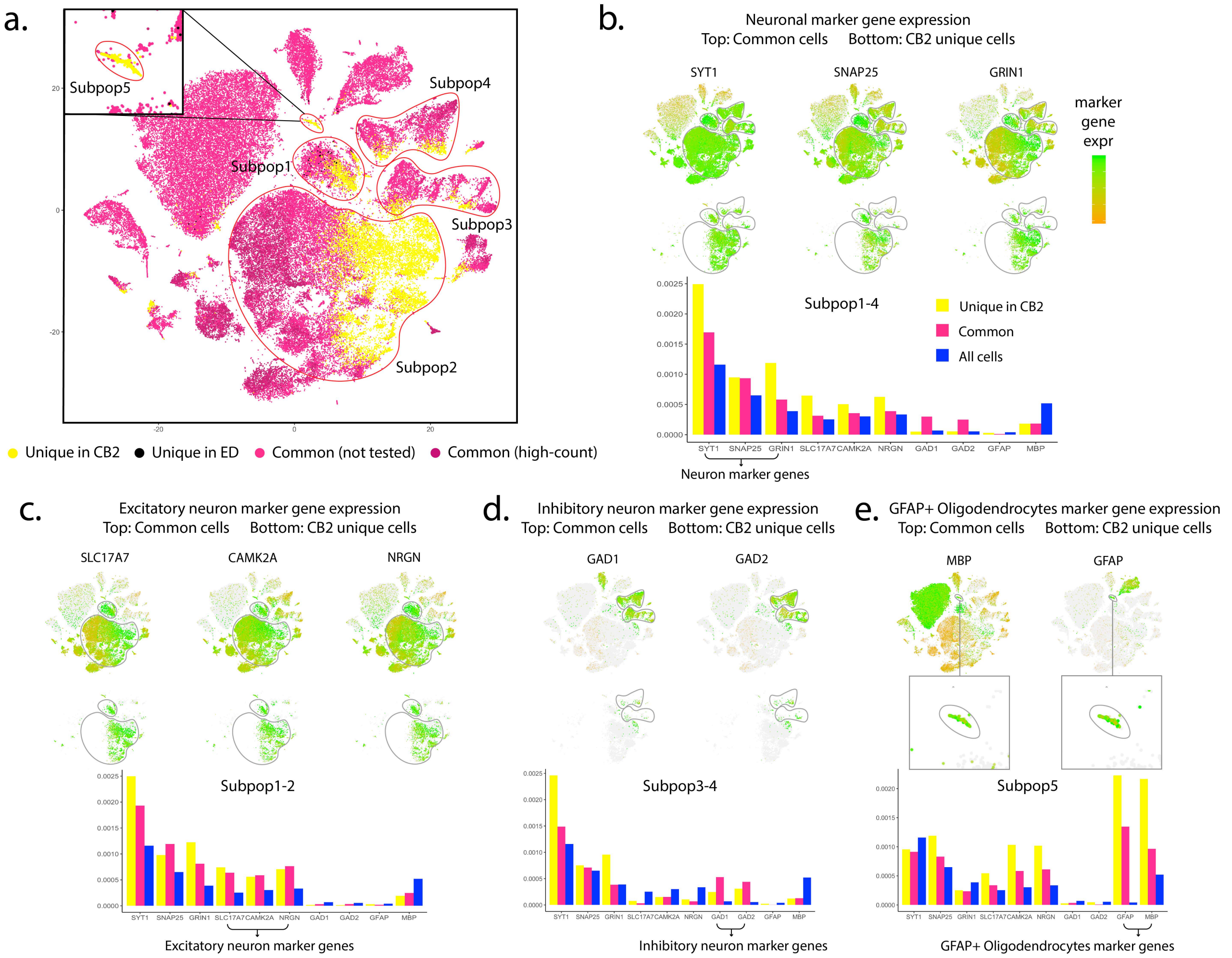


**Supplementary Figure 6**: Additional analysis of Alzheimer dataset. Mathys *et al*. 2019[3] considered marker genes to identify neurons (SYT1, SNAP25, GRIN1), excitatory neurons (SLC17A7, CAMK2A, NRGN), inhibitory neurons (GAD1, GAD2) and oligodendrocytes (MBP). (a) The same t-SNE plot as in **Figure 2**. Panels (b)-(e) show that t-SNE plot colored by marker gene expression in all cells (upper) as well as those identified uniquely by CB2 (lower). Distribution plots of the 10 marker genes within the specified subpopulations are shown in the bottom panels for the common cells (pink) and those uniquely identified by CB2 (yellow). Expression levels of these markers calculated across all cells are shown in blue as a reference. Cells uniquely identified by CB2 have marker gene expression patterns similar to those observed in the cells identified in common between CB2 and ED, providing further support that they are real cells. The novel subpopulation identified by CB2 is the only subpopulation showing high expression of both oligodendrocyte and astrocyte marker genes, suggesting that this group may be mixed phenotype glial cells[4] (GFAP+ oligodendrocytes).

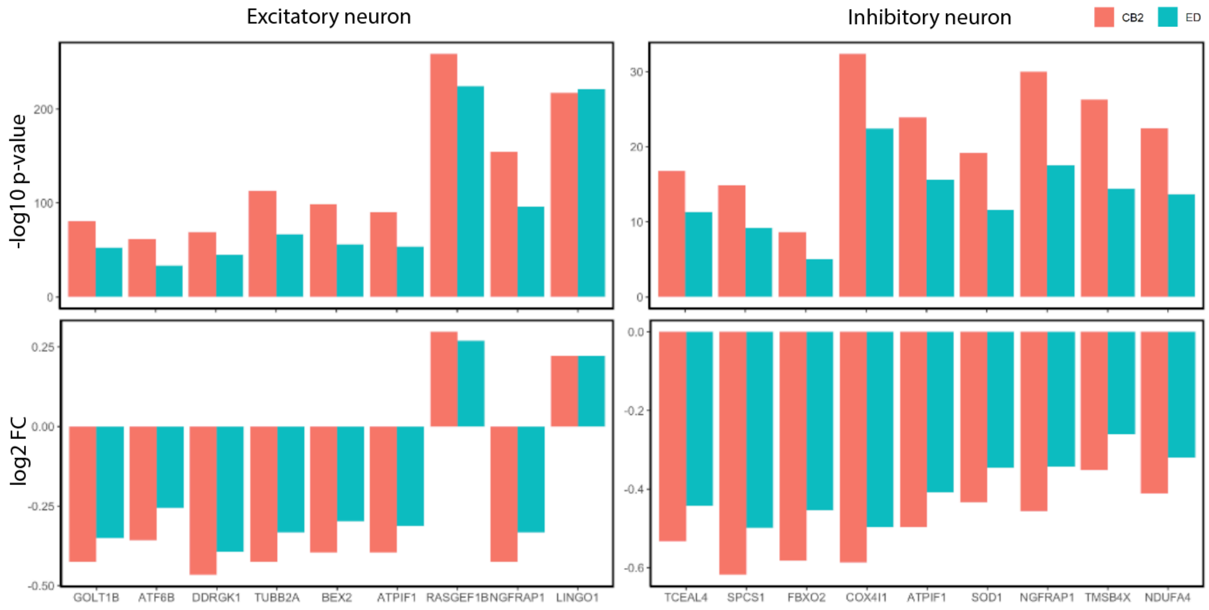


**Supplementary Figure 7**: Differential Expression analysis between Alzheimer's disease (AD) cases and controls was conducted using cells identified by CB2 (salmon) and ED (turquoise). Shown in the upper panels are the -log10 p-values for 9 genes known to be differentially expressed between AD-pathology and control cells in excitatory neurons (left: GOLT1B, ATF6B, DDRGK1, TUBB2A, BEX2, ATPIF1, RASGEF1B, NGFRAP1, LINGO1) and 9 genes known to be differentially expressed in inhibitory neurons (right: TCEAL4, SPCS1, FBXO2, COX4I1, ATPIF1, SOD1, NGFRAP1, TMSB4X, NDUFA4)[3]. Log2 fold changes of the mean expression in AD-pathology vs. control cells are also shown (lower panels) for each gene in each cell type. CB2 improves downstream DE analysis by showing more significant p-values and stronger fold changes.

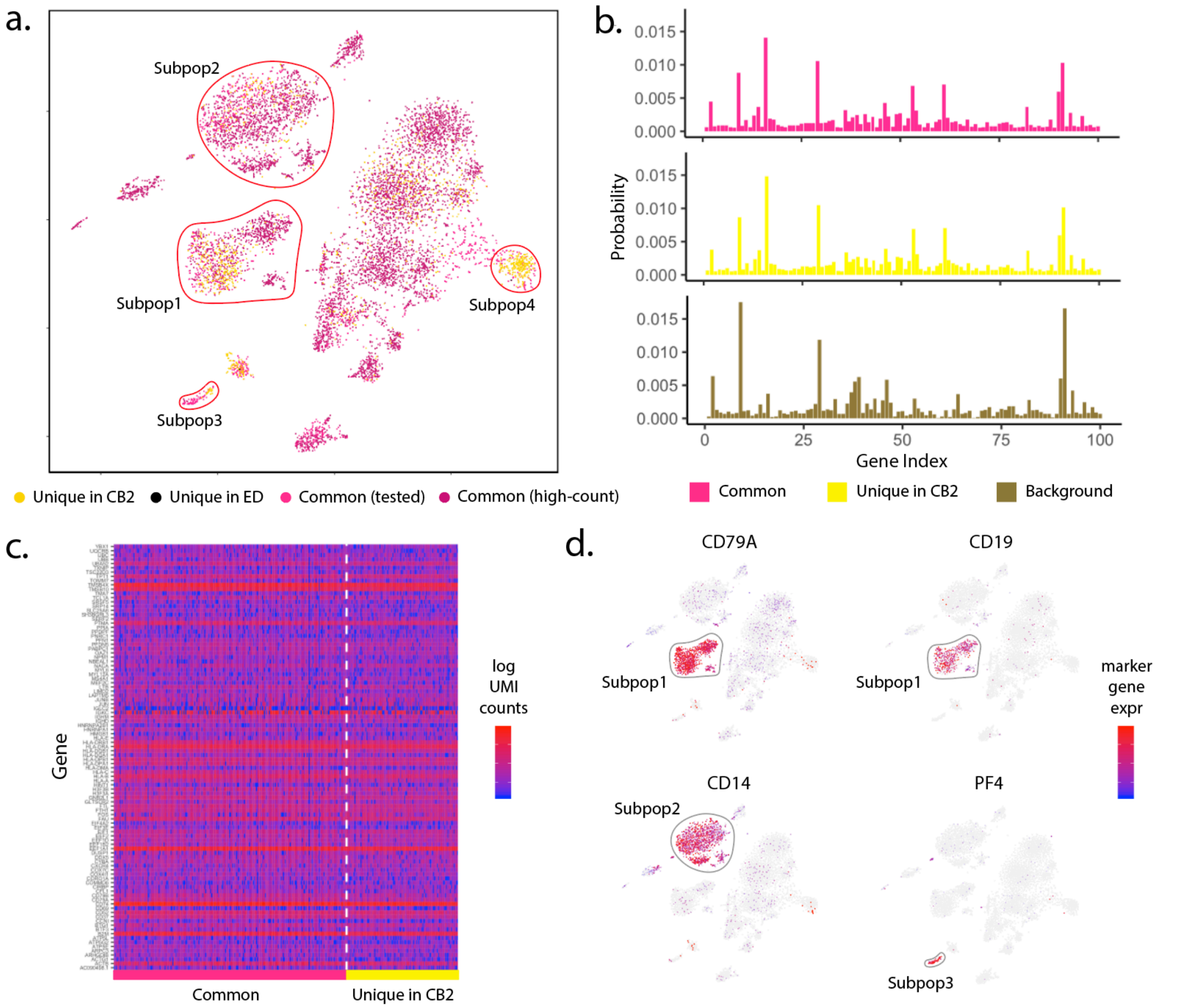


**Supplementary Figure 8**: Analysis of PBMC8K dataset. (a) t-SNE plot of cells identified by CB2 and ED. High-count barcodes exceeding an upper threshold are identified as real cells by both methods without a statistical test (dark pink); barcodes identified as cells by both methods following statistical test are shown in pink. Cells identified uniquely by CB2 (yellow) and ED (black) are also shown. (b) Distribution plots of the 100 genes having highest average expression in Subpop1 are shown for cells identified by both CB2 and ED (upper) and identified uniquely by CB2 (middle). The estimated background distribution is also shown (lower). Cells uniquely identified by CB2 in Subpop1 have a distribution similar to other Subpop1 cells and differ from the background. (c) Heatmap of log transformed raw UMI counts for the same 100 genes for barcodes identified by CB2 and ED (left) and barcodes uniquely identified by CB2 (right). (d) t-SNE plots of cells colored by known marker genes for B-cells (CD79A, CD19), CD14+ Monocytes (CD14), and Megakaryocytes (PF4)[1].

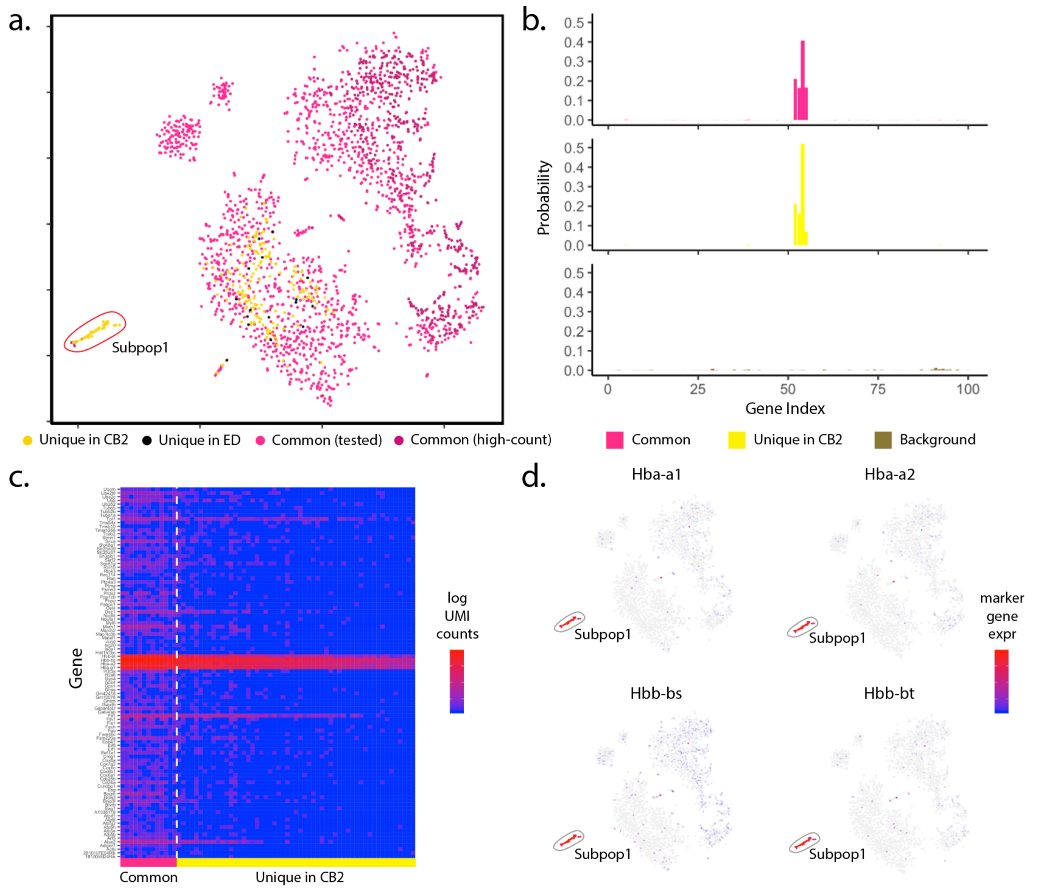


**Supplementary Figure 9**: Analysis of mbrain1K dataset. (a) t-SNE plot of cells identified by CB2 and ED. High-count barcodes exceeding an upper threshold are identified as real cells by both methods without a statistical test (dark pink); barcodes identified as cells by both methods following statistical test are shown in pink. Cells identified uniquely by CB2 (yellow) and ED (black) are also shown. (b) Distribution plots of the 100 genes having highest average expression in Subpop1 are shown for cells identified by both CB2 and ED (upper) and identified uniquely by CB2 (middle). The estimated background distribution is also shown (lower). Cells uniquely identified by CB2 in Subpop1 have a distribution similar to other Subpop1 cells and differ from the background. (c) Heatmap of log transformed raw UMI counts for the same 100 genes for barcodes identified by CB2 and ED (left) and barcodes uniquely identified by CB2 (right). (d) t-SNE plots of cells colored by expression of hemoglobin-related marker genes (Hba-a1, Hba-a2, Hbb-bs, Hbb-bt). (b)-(d) indicate that CB2 reveals a novel subpopulation of cells with expression patterns consistent with high stress and neurogenerative disorders in both human[5] and mouse[6] brain studies.

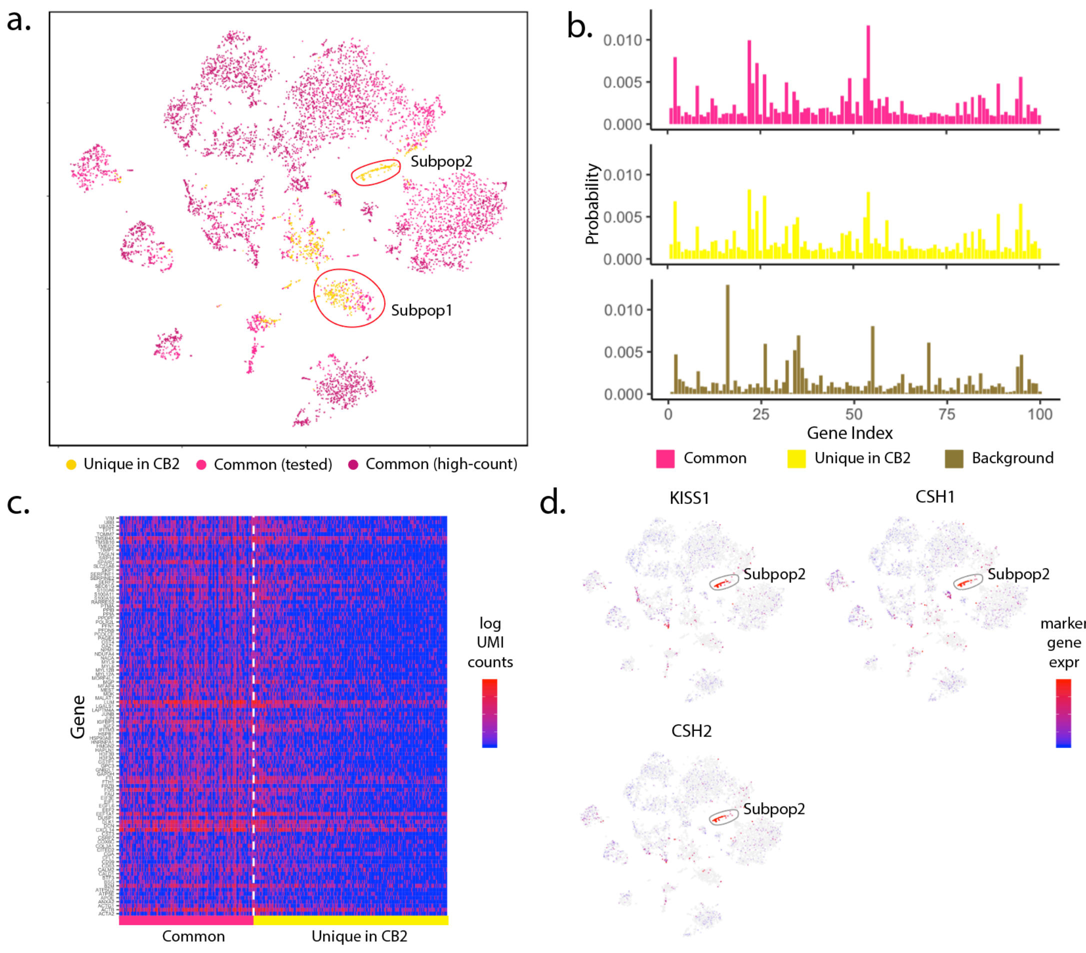


**Supplementary Figure 10**: Analysis of placenta dataset. (a) t-SNE plot of cells identified by CB2 and ED. High-count barcodes exceeding an upper threshold are identified as real cells by both methods without a statistical test (dark pink); barcodes identified as cells by both methods following statistical test are shown in pink. Cells identified uniquely by CB2 (yellow) and ED (black) are also shown. (b) Distribution plots of the 100 genes having highest average expression in Subpop1 are shown for cells identified by both CB2 and ED (upper) and identified uniquely by CB2 (middle). The estimated background distribution is also shown (lower). Cells uniquely identified by CB2 in Subpop1 have a distribution similar to other Subpop1 cells and differ from the background. (c) Heatmap of log transformed raw UMI counts for the same 100 genes for barcodes identified by CB2 and ED (left) and barcodes uniquely identified by CB2 (right). (d) t-SNE plots of cells colored by expression of a placenta-specific marker gene (KISS1) and marker genes related to placental lactogen secretion (CSH1, CSH2)[7]. Panels (a) and (d) indicate that CB2 reveals a novel subpopulation (Subpop2) that may be related to placental lactogen secretion.

| Dataset | High-count cells (untested) | Tested cells identified by CB2 and ED | CB2 unique cells | ED unique cells | CB2 unique cells in existing subpopulation | CB2 unique cells in novel subpopulation |
|---|---|---|---|---|---|---|
| Alzheimer | 12143 | 57278 | 10689 / 57278 (18.66%) | 50 / 57278 (0.09%) | 6819 / 10689 (63.79%) | 3870 / 10689 (36.21%) |
| PBMC8K | 6708 | 1445 | 1165 / 1445 (80.62%) | 2 / 1445 (0.14%) | 1165 / 1165 (100%) | 0 / 1165 (0%) |
| PBMC33K | 23491 | 11762 | 424 / 11762 (3.60%) | 0 / 11762 (0%) | 424 / 424 (100%) | 0 / 424 (0%) |
| mbrain1K | 581 | 1469 | 221 / 1469 (15.04%) | 16 / 1469 (1.09%) | 166 / 221 (75.11%) | 55 / 221 (24.89%) |
| mbrain9K | 6048 | 5685 | 1265 / 5685 (22.25%) | 98 / 5685 (1.72%) | 1057 / 1265 (83.56%) | 208 / 1265 (16.44%) |
| PanT4K | 3398 | 1700 | 261 / 1700 (15.35%) | 0 / 1700 (0%) | 261/ 261 (100%) | 0 / 261 (0%) |
| MALT | 3378 | 981 | 494 / 981 (50.36%) | 2 / 981 (0.20%) | 216 / 494 (43.72%) | 278 / 494 (56.28%) |
| PBMC4K | 2145 | 6516 | 1003 / 6516 (15.39%) | 0 / 6516 (0%) | 1003 / 1003 (100%) | 0 / 1003 (0%) |
| jurkat | 2565 | 953 | 175 / 953 (18.36%) | 0 / 953 (0%) | 175 / 175 (100%) | 0 / 175 (0%) |
| T293 | 2299 | 797 | 48 / 797 (6.02%) | 2 / 797 (0.25%) | 48 / 48 (100%) | 0 / 48 (0%) |
| placenta | 4349 | 2947 | 637 / 2947 (21.62%) | 1 / 2947 (0.03%) | 637 / 637 (100%) | 0 / 637 (0%) |

**Supplementary Table 1:** The number of cells identified by CB2, ED, or both in 11 case study datasets.

| Dataset | Link |
| --- | --- |
| Alzheimer | https://www.synapse.org/#!Synapse:syn16780177 |
| PBMC8K | https://support.10xgenomics.com/single-cell-gene-expression/datasets/2.1.0/pbmc8k |
| PBMC33K | https://support.10xgenomics.com/single-cell-gene-expression/datasets/1.1.0/pbmc33k |
| mbrain1K | https://support.10xgenomics.com/single-cell-gene-expression/datasets/2.1.0/neurons_900 |
| mbrain9K | https://support.10xgenomics.com/single-cell-gene-expression/datasets/2.1.0/neuron_9k |
| PanT4K | https://support.10xgenomics.com/single-cell-gene-expression/datasets/2.1.0/t_4k |
| MALT | https://support.10xgenomics.com/single-cell-gene-expression/datasets/3.0.0/malt_10k_protein_v3 |
| PBMC4K | https://support.10xgenomics.com/single-cell-gene-expression/datasets/2.1.0/pbmc4k |
| jurkat | https://support.10xgenomics.com/single-cell-gene-expression/datasets/1.1.0/jurkat |
| T293 | https://support.10xgenomics.com/single-cell-gene-expression/datasets/1.1.0/293t |
| placenta | https://jmlab-gitlab.cruk.cam.ac.uk/publications/EmptyDrops2017-DataFiles |

**Supplementary Table 2**: Links to all datasets used in this study.

| Dataset | Threshold | | |
| --- | --- | --- | --- |
| | **90%** | **80%** | **70%** |
| Alzheimer | 1 | 2 | 2 |
| PBMC8K | 0 | 0 | 0 |
| PBMC33K | 0 | 0 | 0 |
| mbrain1K | 0 | 1 | 1 |
| mbrain9K | 0 | 1 | 1 |
| PanT4K | 0 | 0 | 0 |
| MALT | 1 | 1 | 2 |
| PBMC4K | 0 | 0 | 0 |
| jurkat | 0 | 0 | 1 |
| T293 | 0 | 0 | 0 |
| placenta | 0 | 0 | 1 |

**Supplementary Table 3**: Number of novel subpopulations identified by CB2 in each dataset.
